# The c-Src inhibitor eCF506 diminishes opioid tolerance creating bias against β-arrestin2 recruitment

**DOI:** 10.1101/2025.07.01.662507

**Authors:** Samuel Singleton, Fraser Nunn, Erika Lace, Álvaro Lorente-Macías, Rachel Toth, Asier Unciti-Broceta, Tim G Hales

**Affiliations:** Institute of Academic Anaesthesia, Division of Systems Medicine, School of Medicine, Ninewells Hospital, University of Dundee, Dundee, DD1 9SY, UK; Edinburgh Cancer Research, CRUK Scotland Centre, Institute of Genetics and Cancer, University of Edinburgh, Edinburgh, EH4 2XR, UK; MRC PPU Reagents and Services, MRC Protein Phosphorylation and Ubiquitylation Unit, School of Life Sciences, Wellcome Trust Biocentre, University of Dundee, Dow Street, Dundee, DD1 5EH, UK

**Keywords:** Analgesia, NXP900, arrestins, tolerance, pain

## Abstract

Opioids reduce severe pain, but persistent use is compromised by tolerance, attenuated by either β-arrestin2 depletion, prompting development of biased opioids limited by partial efficacy, or c-Src kinase inhibitors, potentially acting through off-target effects. We tested eCF506, a conformationally selective c-Src inhibitor, on morphine antinociception and examined its effect on μ receptor signaling and β-arrestin2 recruitment. Oral eCF506 inhibited morphine tolerance in C57BL/6J mice. Exposure of PathHunter CHO cells to eCF506 did not affect inhibition of cAMP accumulation by the μ agonist, DAMGO, but reduced β-arrestin2 recruitment. This effect, mimicked by targeted degradation of c-Src, occurred through inhibition of c-Src catalytic function as evidenced by its diminution by the catalytically inactive Src250-536(K298M) construct. This mutant also restricted the effect of c-Src inhibitors on β-arrestin2 recruitment. eCF506 additionally increased surface expression of μ receptors and limited their internalization by endomorphin-2 but did not alter DAMGO-evoked GRK-mediated receptor phosphorylation. These findings suggest that eCF506 prolongs opioid antinociception by inducing signalling bias, diminishing β-arrestin2-mediated μ receptor regulation.

## Introduction

Chronic pain, pain persisting for 3 months or more, affects one third of the population and is the leading cause of years lived with disability, highlighting the need for effective analgesic strategies (Cohen et al., 2021).

Opioid analgesics activate μ receptors to provide powerful relief from moderate to severe acute pain by signaling through Gα_i/o_ proteins to inhibit voltage activated Ca^2+^ channels (VACCs) and activate G-protein activated inwardly rectifying K^+^ (GIRK) channels and suppress neuronal excitation (Matthes et al., 1996; Heinke et al., 2011). However, their use for managing chronic pain is compromised by the development of analgesic tolerance and hyperalgesia (Colvin et al., 2019). In mouse models morphine tolerance and associated hypersensitivity require μ receptors, β-arrestin2 and the non-receptor tyrosine kinase, c-Src (Bohn et al., 2000; Bull et al., 2017; Singleton and Hales, 2023).

Activated μ receptors are targeted for G protein receptor kinase (GRK) mediated c-terminal phosphorylation that enhances the affinity for β-arrestin2 and initiates the processes of desensitisation and internalisation with the involvement of c-Src (Walwyn et al., 2007; Williams et al., 2013). Strategies that diminish the influence of β-arrestin2 and/or c-Src on μ receptors may prolong inhibitory coupling to provide sustained analgesia and improve the long-term use of opioids. Indeed, either a lack of β-arrestin2 or pharmacological inhibition of c-Src in wild type mice enhances μ receptor coupling to VACCs and attenuates morphine-evoked antinociceptive tolerance (Bohn et al., 2000; Bull et al., 2017; Walwyn et al., 2007).

The identification of opioids that exhibit bias in favour of G protein signaling and against β-arrestin2 recruitment is an approach that has received considerable attention (Kelly et al., 2023). However, such agonists tend to have low efficacy complicating their characterisation as truly biased. Inhibitors of c-Src offer an alternative approach to diminish β-arrestin2 mediated tolerance and opioid hypersensitivity (Bull et al., 2017; Singleton and Hales, 2023).

However, prior work has utilised the promiscuous Src family kinase inhibitor, PP2, or the clinically used non-selective c-Abl/c-Src inhibitor dasatinib, which can cause off-target effects, including cardiac events through inhibition of c-Abl (Steegmann et al., 2012). These limitations have the potential to confound interpretations for the role of c-Src in μ receptor signal transduction and diminish enthusiasm for using dasatinib as an adjunct to improve opioid analgesia. Approaches with enhanced selectivity for targeting c-Src kinase may be leveraged to modulate opioid signal transduction and provide sustained analgesia by μ receptor agonists.

In this study, we assessed the impact of conformationally selective c-Src kinase inhibition using eCF506 (a.k.a. NXP900), a drug in clinical development that improves selectivity and tolerability by binding c-Src in its autoinhibited conformation (Fraser et al., 2016; Temps et al., 2021). We examined the impact of eCF506 on the development of antinociceptive tolerance to morphine and establish the contributions of c-Src catalytic and non-catalytic domains on the recruitment of β-arrestin2 to μ receptors in *vitro* using constructs lacking either one or both domains (Miller et al., 2000).

## Results

### eCF506 delays the development of analgesic tolerance to morphine

Tolerance to the analgesic effect of morphine in an assay of thermal nociception is inhibited in mice administered the Src-family kinase inhibitors dasatinib or PP2 (Bull et al., 2017). We assessed the

development of analgesic tolerance to once-daily injections with morphine in mice administered the conformationally selective c-Src inhibitor, eCF506 (Fraser et al., 2016; Temps et al., 2021).

Administration of eCF506 did not affect basal tail withdrawal latency (TWL) or morphine antinociception in WT mice (Supplementary Figure 1A-B). Mean basal TWL after administration of vehicle was 4.3 ± 0.4 s, while after administration of eCF506 (20 or 80 mg/Kg) values were 4.8 ± 0.6 s and 4.4 ± 0.5 s, respectively (Supplementary Figure 1A). The mean antinociceptive potencies (ED_50_) of morphine in the tail withdrawal assay following administration of vehicle, eCF506 (20 mg/Kg) or eCF506 (80 mg/Kg) were: 1.5 [0.6 – 2.5], 1.5 [0.8 – 2.2] and 1.0 [0.5 – 1.4] mg/Kg, respectively (Supplementary Figure 1B).

We continued to assess morphine antinociception for 10 consecutive days following once-daily administration of vehicle or eCF506 (20 or 80 mg/Kg) 30 minutes prior to morphine (10 mg/Kg) to assess the impact of c-Src inhibition by eCF506 on the development of antinociceptive tolerance. Morphine caused the development of antinociceptive tolerance in wild type mice administered with vehicle as revealed by a gradual decline in the maximum possible effect (MPE) from day 7 (Figure 1A), and a reduction in the ED_50_ of morphine to 9.7 [7.8 – 11.5] mg/Kg on day 10 (Supplementary Figure 1C). By contrast, the decline in %MPE was delayed until day 9 and morphine’s ED_50_ was reduced to 4.2 [1.1 – 7.3] mg/Kg in mice administered with eCF506 (20 mg/Kg). There was no decline in morphine antinociception (% MPE) across the 10 days in mice receiving eCF506 (80 mg/Kg) and the ED_50_ on day 10 was 2.6 [1.7 – 3.6] mg/Kg (Figure 8A; Supplementary Figure 1D-E).

**Figure 1:**
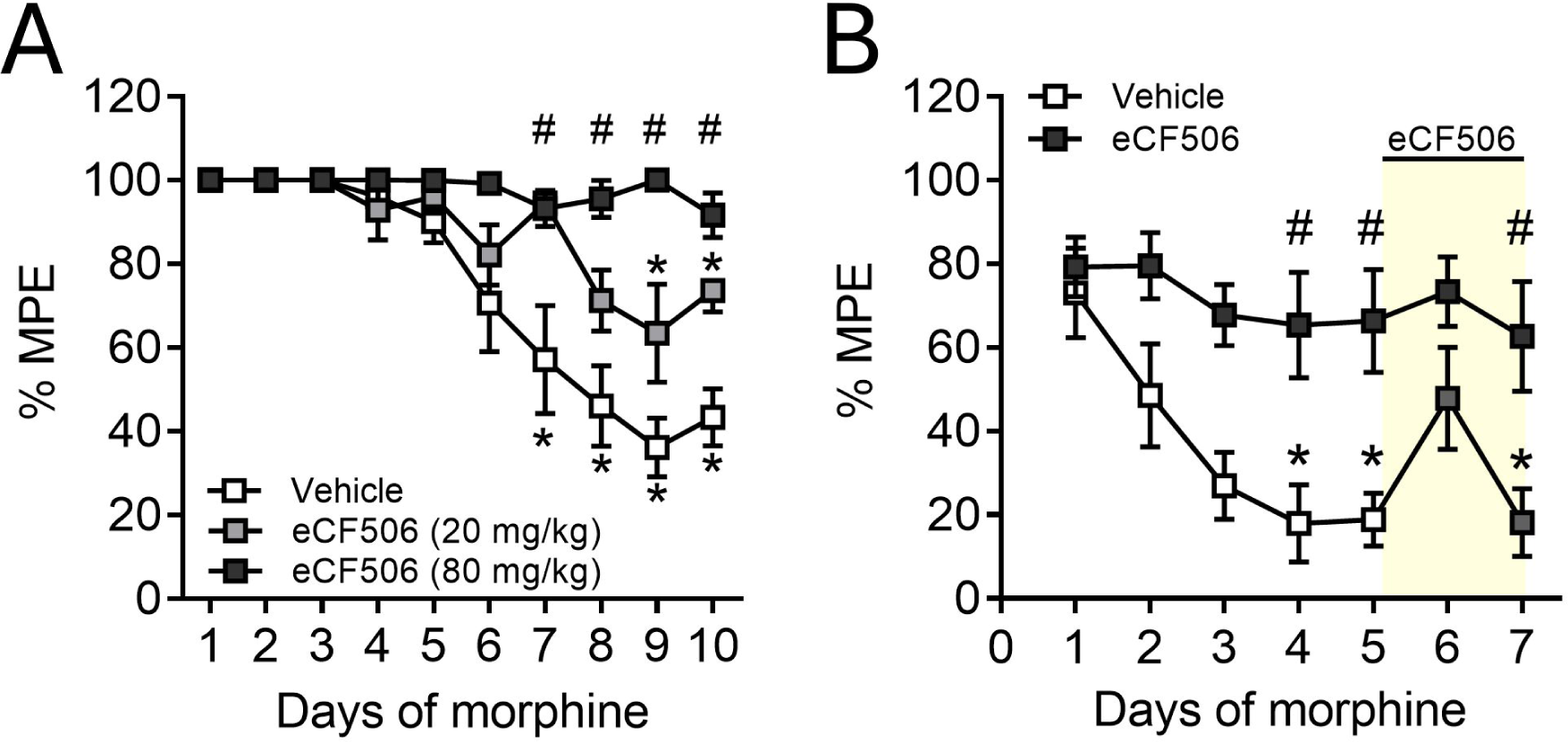
Inhibition of c-Src using eCF506 prevents the development of morphine analgesic tolerance. Maximum possible effect (%MPE; see Methods) of morphine (10 mg/kg) administered once-daily (s.c.) to prolong tail withdrawal latencies from warm (48°C) water in male and female (A) WT or (B) μ+/- C57BL/6J mice. (**A**) WT mice were administered with vehicle (0.9% normal saline with 2% DMSO and 2% Kolliphor EL) or eCF506 (20 - 80 mg/kg reconstituted in vehicle) and tail withdrawal latencies assessed prior to, and 30 minutes after, injection with morphine, once-daily for 10 consecutive days. (**B**) Vehicle or eCF506 (80 mg/kg) was administered to μ+/- mice and tail withdrawal latencies assessed once-daily using the same approach for 7 consecutive days. On days 6 and 7, all μ+/- mice were administered with eCF506 (80 mg/kg) to assess reversal of established morphine antinociceptive tolerance (yellow shaded region). Dose-response relationships of morphine antinociception in WT and μ+/- mice on the first and last day of injections are shown in Supplementary Figure 1. Data are the mean ± SEM of (A) 7 (vehicle) and n = 8 (eCF506) mice and (B) n = 6 mice (vehicle, eCF506 20 mg/kg and eCF506 80 mg/kg).

The development of antinociceptive tolerance to morphine is accelerated and more profound in μ+/- mice expressing 50% fewer receptors (Sora et al., 2001; Bull et al., 2017). We determined first, whether c-Src inhibition by eCF506 (80 mg/Kg) could provide sustained antinociception even in μ+/- mice and then, whether established tolerance could be reversed by the Src inhibitor (Figure 1B). The potency of morphine to cause antinociception in μ+/- mice (mean ED_50_ 4.4 [3.5 – 5.5] mg/Kg) was reduced when compared to WT mice (Supplementary Figure 8B), consistent with reduced μ receptor expression. Morphine (10 mg/Kg) antinociception rapidly declined by day 4 in μ+/- mice receiving vehicle whereas antinociception was sustained in μ+/- mice receiving eCF506 (Figure 1B; Supplementary Figure 1F). On days 6 and 7 of morphine injections, we administered eCF506 (80 mg/Kg) to all mice to determine whether c-Src inhibition could reverse established tolerance (Figure 1B). On day 6, administration of eCF506 caused transient restoration of morphine antinociception in μ+/- mice previously administered with vehicle and rendered tolerant through daily morphine (Figure 1B). However, by day 7, this effect diminished, and morphine failed to cause antinociception. By contrast, morphine continued to cause antinociception in mice that received daily oral eCF506 from day 1.

Together, these data demonstrate that inhibition of c-Src using eCF506 prevents the development of antinociceptive tolerance when administered as an adjunct to morphine and causes a transient

reversal of established tolerance.

### Phosphorylation of c-Src kinase by μ receptor agonists correlates with agonist efficacy for recruiting β-arrestin2

We examined the effect of DAMGO, fentanyl, morphine, oxycodone, buprenorphine and TRV130 on β-arrestin2 recruitment to μ receptors using PathHunter CHO cells stably overexpressing human μ receptors (Figure 2A). Mean efficacy (E_MAX_) and potency (pEC_50_) values are summarised in Supplementary Table 1. A one-way ANOVA with Bonferroni corrections revealed that all agonists had partial efficacy at recruiting β-arrestin2 to μ receptors when compared to DAMGO (F5,33 = 80.4). By contrast, only oxycodone was significantly less potent than DAMGO for recruiting β-arrestin2 (Supplementary Table 1).

**Figure 2:**
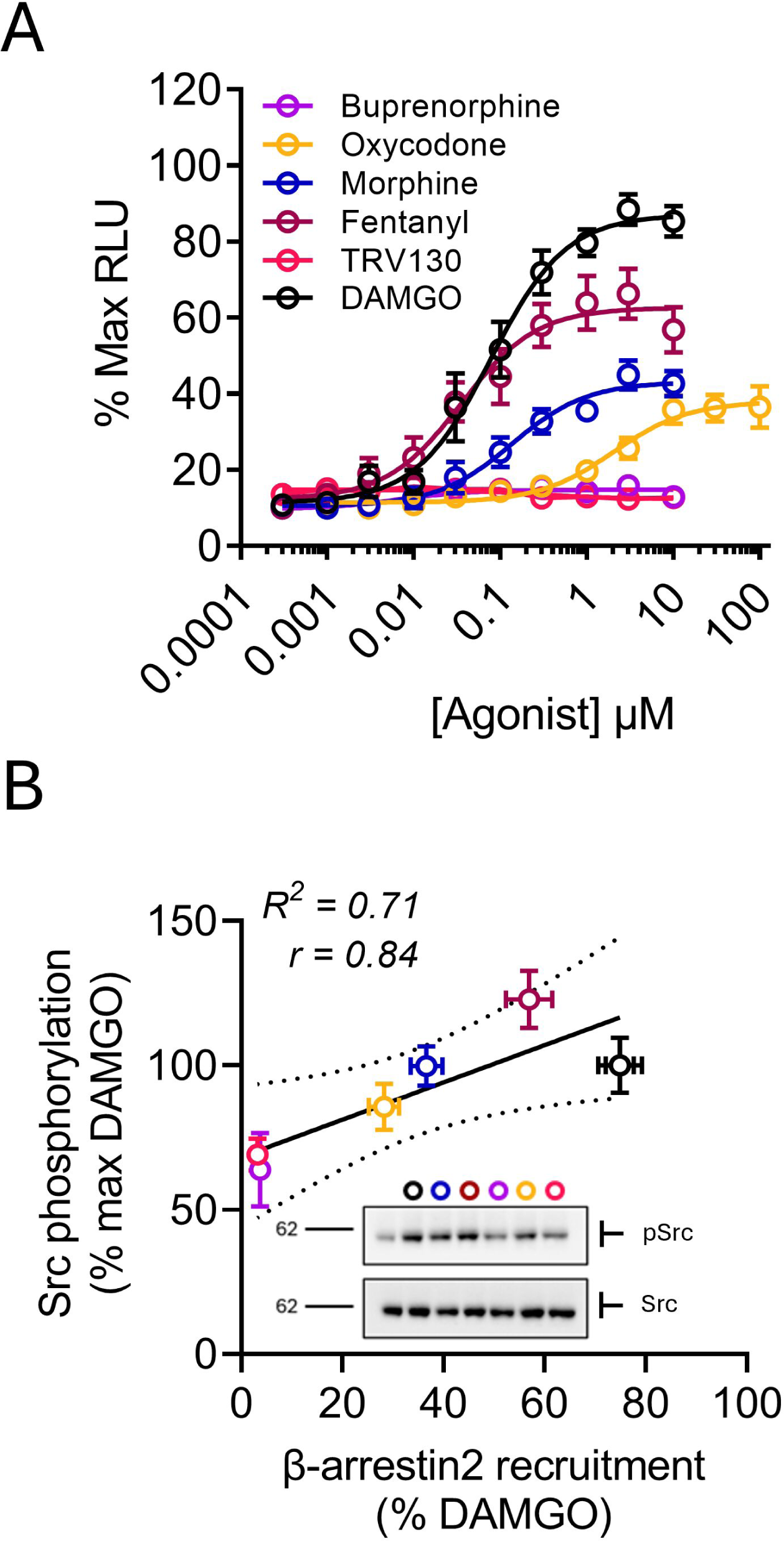
Recruitment of β-arrestin2 and c-Src Phosphorylation by μ receptor agonists. (**A**) Concentration-response relationships of DAMGO, morphine, fentanyl, oxycodone, buprenorphine and TRV130 to recruit β-arrestin2 to human μ receptors in PathHunter CHO cells. Data are expressed as a percentage of maximum luminescence (% Max RLU) produced by DAMGO present in duplicate on each plate. Data from individual replicates were plotted and fitted with logistics functions to derive efficacy (E_MAX_) and potency (EC_50_) parameters, presented in Supplementary Table 1. (**B**) Correlation between agonist E_MAX_ for recruiting β-arrestin2 (A) and levels of c-Src phosphorylation at Y416 assessed in PathHunter CHO cells by western blot following 15-minute exposure to saturating (10 μM) concentrations of DAMGO, morphine, fentanyl, oxycodone, buprenorphine or TRV130 (Supplementary Figure 2A-B). Phosphorylation is expressed relative to total c-Src present in each cell lysate and as percentage of maximum phosphorylation caused by DAMGO (Supplementary Figure 2A). Data are the mean ± SEM of 7 replicates (A) and 4 replicates (B).

β-arrestin2 is required to target c-Src to μ receptors as revealed by a lack of μ receptor agonist evoked phosphorylation of c-Src in dorsal root ganglia of β-arrestin2-/- mice (Walwyn et al., 2007). Therefore, we assessed whether these agonists were also capable of activating c-Src by measuring the phosphorylation of residue Y416, a hallmark of c-Src activation (Roskoski, 2005), after a 15-minute exposure to agonists, using western blotting (Figure 2B; Supplementary Figure 2). DAMGO, fentanyl, morphine, oxycodone, buprenorphine and TRV130 (10 μM) all caused c-Src activation as revealed by an enhanced ratio of phosphorylated (Y416) to total c-Src when compared to PathHunter CHO cells exposed to DMSO (Supplementary Figure 2).

Correlating the efficacy of each agonist to recruit β-arrestin2 against their efficacy to phosphorylate c-Src revealed a Pearson correlation coefficient (r) of 0.84 (95% CIs: 0.10 - 0.98). These data demonstrate that μ receptor agonists activate c-Src kinase, with phosphorylation at residue Y416 correlating to agonist efficacy for β-arrestin2 recruitment.

### Inhibition of c-Src kinase limits β-arrestin2 recruitment to μ receptors without affecting G-protein activation

μ Receptors can be phosphorylated by c-Src in mouse embryonic fibroblasts depleted of β-arrestins (Zhang et al., 2009; Zhang et al., 2017) implying that c-Src may regulate signalling independent of β- arrestin2 binding. Therefore, we determined the impact of inhibiting c-Src kinase using eCF506 or PP2 on the concentration-response relationships of DAMGO to recruit β-arrestin2 to μ receptors (Figure 3A) and inhibit the accumulation of cAMP (Figure 3B). PathHunter CHO cells were either exposed to 1 μM eCF506 or PP2, or to PP3, a c-Src inactive analogue to PP2, overnight (16-hours) and concentration-response relationships of DAMGO were compared to that of cells exposed to an equal volume of DMSO. Mean efficacy (E_MAX_) and potency (pEC_50_ or pIC_50_) values are summarised in Table 1.

**Figure 3:**
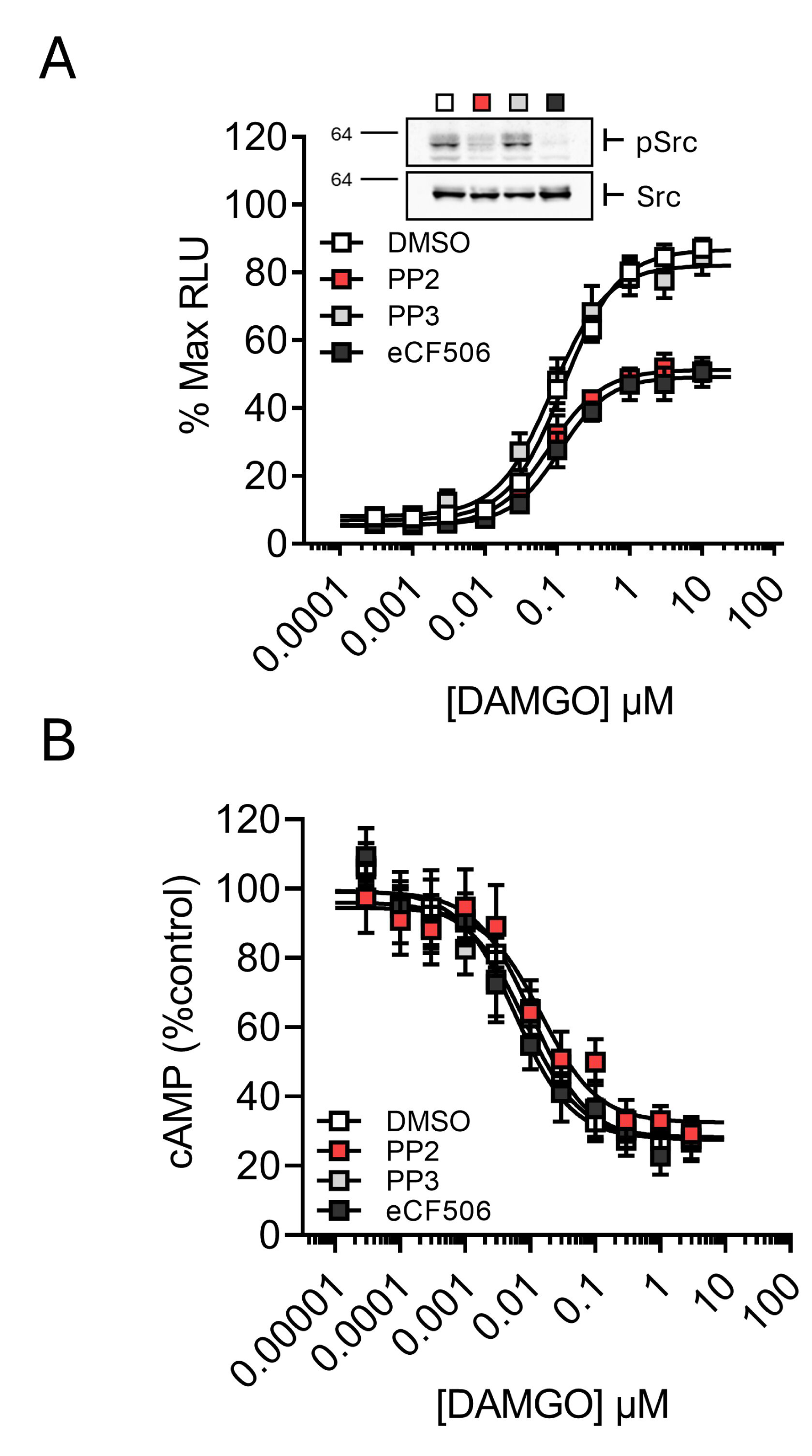
Inhibition of c-Src reduces β-arrestin2 recruitment to μ receptors. Concentration-response relationships of (**A**) DAMGO as a recruiter of β-arrestin2 to human μ receptors in PathHunter CHO cells and (**B**) DAMGO as an inhibitor of forskolin (30 μM) stimulated cAMP accumulation in the same cell line transiently expressing the pGloSensor-22F protein following 16 h exposure to 1 μM PP2, PP3, eCF506 or an equal volume of DMSO. (A) Data are expressed as a percentage of maximum luminescence (% Max RLU) produced by DAMGO in DMSO exposed cells present in duplicate on each 96-well plate. The insert demonstrates c-Src phosphorylation at Y416 (upper panel) and total c-Src (lower panel) expression assessed using western blot in each cell lysate following overnight exposure and confirms inhibition of c-Src phosphorylation. (B) Data are expressed as a percentage of luminescence in each well measured prior to the addition of agonist (% control). Data from individual replicates were plotted and fitted with logistics functions to derive efficacy (E_MAX_) and potency (EC_50_/IC_50_) parameters, presented in Table 1. Data are the mean ± SEM of 5 replicates (A) and 8 replicates (B).

**Table 1:** Efficacies and potencies of DAMGO to recruit β-arrestin2 and inhibit the accumulation of cAMP following overnight c-Src inhibition. Individual concentration-response relationships of DAMGO to recruit β-arrestin2 (Figure 3A) or inhibit cAMP accumulation in PathHunter CHO cells with full (Figure 3B) or partial (Supplementary Figure 3E-H) μ receptor availability were fitted with a logistics function to yield efficacy and potency values. A one-way ANOVA with Dunnett pairwise comparisons revealed that the E_MAX_ of DAMGO to recruit β-arrestin2 was reduced by PP2 and eCF506 when compared to DMSO whereas the E_MAX_ of DAMGO following exposure to PP3 was similar to DMSO (F3,16 = 33.4). By contrast, DAMGO was equally potent in all cells (F3,16 = 1.8). In the cAMP assay, DAMGO achieved a similar E_MAX_ (D3,28 = 0.8) and had the same apparent potency (F3,28 = 1.3) in all cells regardless of overnight treatment. Exposure to β-FNA (100 nM) to restrict μ receptor availability reduced the potency of DAMGO compared to its potency with full μ receptor receptor availability (p < 0.05) but this was not influenced by c-Src inhibition. Limited μ receptor availability also did not change the E_MAX_ of DAMGO to inhibit cAMP accumulation. * p < 0.05 compared to DAMGO and # p < 0.05 compared to the same agonist with full μ receptor availability.

| | β-arrestin2 | | cAMP | | cAMP + β-FNA (100 nM) | |
| --- | --- | --- | --- | --- | --- | --- |
| | $E_{MAX}$ | $pEC_{50}$ | $E_{MAX}$ | $pIC_{50}$ | $E_{MAX}$ | $pIC_{50}$ |
| DMSO | $80 \pm 2$ | 6.9 [6.8-7.0] | $77 \pm 7$ | 7.9 [7.5-8.3] | $70 \pm 7$ | 7.3 [7.1-7.7]# |
| PP2 | $45 \pm 2^*$ | 7.1 [7.0-7.2] | $64 \pm 4$ | 7.9 [7.6-8.2] | $73 \pm 11$ | 7.3 [6.9-7.4]# |
| PP3 | $74 \pm 4$ | 7.1 [6.9-7.2] | $75 \pm 5$ | 8.2 [7.9-8.2] | $73 \pm 2$ | 7.2 [7.0-7.5]# |
| eCF506 | $43 \pm 4^*$ | 6.9 [6.8-7.1] | $77 \pm 6$ | 8.2 [8.0-8.7] | $74 \pm 2$ | 7.2 [7.0-7.3]# |

Exposure to eCF506 or PP2 diminished the E_MAX_ of DAMGO to recruit β-arrestin2 to the μ receptor (F3,16 = 33.4) without affecting its potency (F3,16 = 1.8) whereas the c-Src inactive analogue, PP3, had no effect (Figure 3A; Table 1). eCF506 (10 nM – 1 μM) also caused a concentration-dependent reduction in the recruitment of β-arrestin2 to μ receptors evoked by DAMGO, morphine, endomorphin-2 and met-enkephalin in PathHunter U2OS cells (Supplementary Figure 3A-D; Supplementary Table 2), demonstrating that c-Src contributes to μ receptor agonist-evoked recruitment of β-arrestin2. By contrast, in the cAMP accumulation assay, neither eCF506 nor PP2 affected the E_MAX_ (F3,28 = 0.8) or apparent potency (F3,28 = 1.3) of DAMGO (Figure 3B; Table 1).

The presence of spare receptors and amplification of G protein signaling through indirect assessment of cAMP levels (Kelly, 2013; Baptista-Hon et al., 2020) compromises straightforward comparisons of the relative efficacies and potencies of DAMGO, which may conceal an effect of c-Src inhibition. Therefore, we additionally compared the concentration-response relationships of DAMGO to inhibit cAMP accumulation whilst restricting the availability of μ receptors using the irreversible antagonist, β-FNA. PathHunter CHO cells were exposed to β-FNA (100 nM) for 1 hour followed by wash off since our prior studies revealed that this concentration eliminates receptor reserve in the cAMP assay (Singleton et al., 2021; Singleton et al., 2024).

The potency of DAMGO to inhibit cAMP accumulation was reduced by prior exposure to β-FNA when compared to cells without prior exposure to β-FNA, consistent with a reduction in available μ receptors (Table 1; Supplementary Figure 3E-H). eCF506 and PP2 similarly had no effect on the E_MAX_ of DAMGO despite limited receptor availability (Table 1). Importantly, the concentrations of eCF506, or PP2 used to inhibit c-Src had no impact on the viability of PathHunter CHO cells when assessed using identical exposure paradigms in the MTT assay (Supplementary Figure 3I). As expected, overnight exposure to eCF506 or PP2 reduced the phosphorylation of c-Src at residue Y416 (Figure 3A; Supplementary Figure 3J). Taken together, these data demonstrate that c-Src inhibition limits the efficacy of μ receptor agonists to recruit β-arrestin2 without affecting their capacity to activate G-proteins.

### Targeted degradation of c-Src kinase limits β-arrestin2 recruitment to μ receptors

To confirm that inhibition of c-Src kinase is responsible for reducing the E_MAX_ of agonists to recruit β- arrestin2 to μ receptors, we assessed the impact of targeted c-Src degradation on DAMGO evoked recruitment of β-arrestin2 to μ receptors using a cereblon-based PROTAC degrader, DAS-5-oCRBN (Mao et al., 2024). PathHunter CHO cells were exposed to DAS-5-oCRBN (1 μM) overnight.

Exposure to 1 μM DAS-5-oCRBN reduced the E_MAX_ of DAMGO to recruit β-arrestin2 to μ receptors to 64 ± 8% compared to cells exposed to DMSO (90 ± 2%). By contrast, the pEC_50_ of DAMGO was 6.4 [6.3 - 6.6] and 6.5 [6.3 – 6.8] with and without exposure to DAS-5-oCRBN, respectively. Degradation was confirmed by western blot using an antibody against total c-Src (Figure 4). Importantly, DAS-5-oCRBN did not affect the viability of PathHunter CHO cells (Supplementary Figure 4A-B). Together, these data demonstrate that a reduction in c-Src expression in PathHunter CHO cells corresponds with limited efficacy of DAMGO to recruit β-arrestin2 to μ receptors.

**Figure 4:**
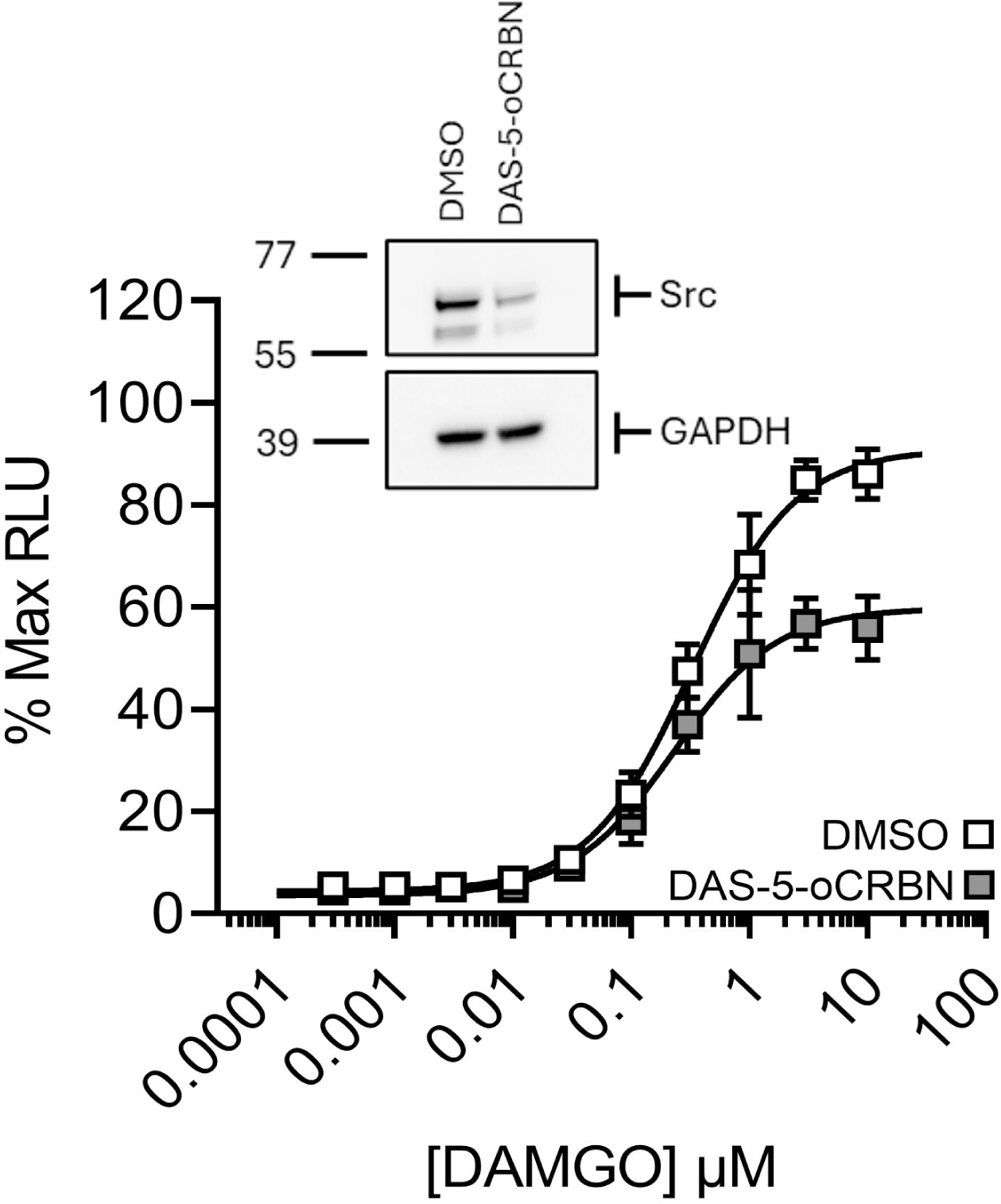
c-Src degradation by DAS-5-oCRBN reduces β-arrestin2 recruitment to μ receptors. Concentration-response relationships of DAMGO to recruit β-arrestin2 to human μ receptors in PathHunter CHO cells following 16-h incubation with escalating concentrations of the selective c-Src degrading PROTAC, DAS-5-oCRBN. Data are expressed as a percentage of maximum luminescence (% Max RLU) produced by DAMGO in DMSO exposed cells present in duplicate on each plate. The insert shows c-Src degradation by DAS-5-oCRBN under identical conditions as confirmed by western blot. Data are the mean ± SEM of 6 replicates.

### Overexpression of mutant c-Src constructs with restricted kinase activity diminish β-arrestin2 recruitment to μ receptors

Having demonstrated that c-Src is required for μ receptor agonists to efficiently recruit β-arrestin2, we next explored the contribution of c-Src catalytic and non-catalytic functions to β-arrestin2 recruitment by comparing the efficacy and potency of DAMGO to recruit β-arrestin2 in PathHunter CHO cells transiently overexpressing WT c-Src (Src1-536) to its efficacy and potency in cells overexpressing truncated c-Src mutants lacking either the protein scaffolding (Src250-536) or kinase domains (Src1-249). The latter retains the capacity to scaffold c-Src substrates (Miller et al., 2000). Mean efficacy (E_MAX_) and potency (pEC_50_) values of DAMGO to recruit β-arrestin2 are summarised in Table 2. Overexpression of each mutant was confirmed via western blot using an antibody recognising the N-terminus of Src or using an HA-tag inserted at the C-terminus in the case of Src1-249 (Figure 5A; Supplementary Figure 5A)

**Figure 5:**
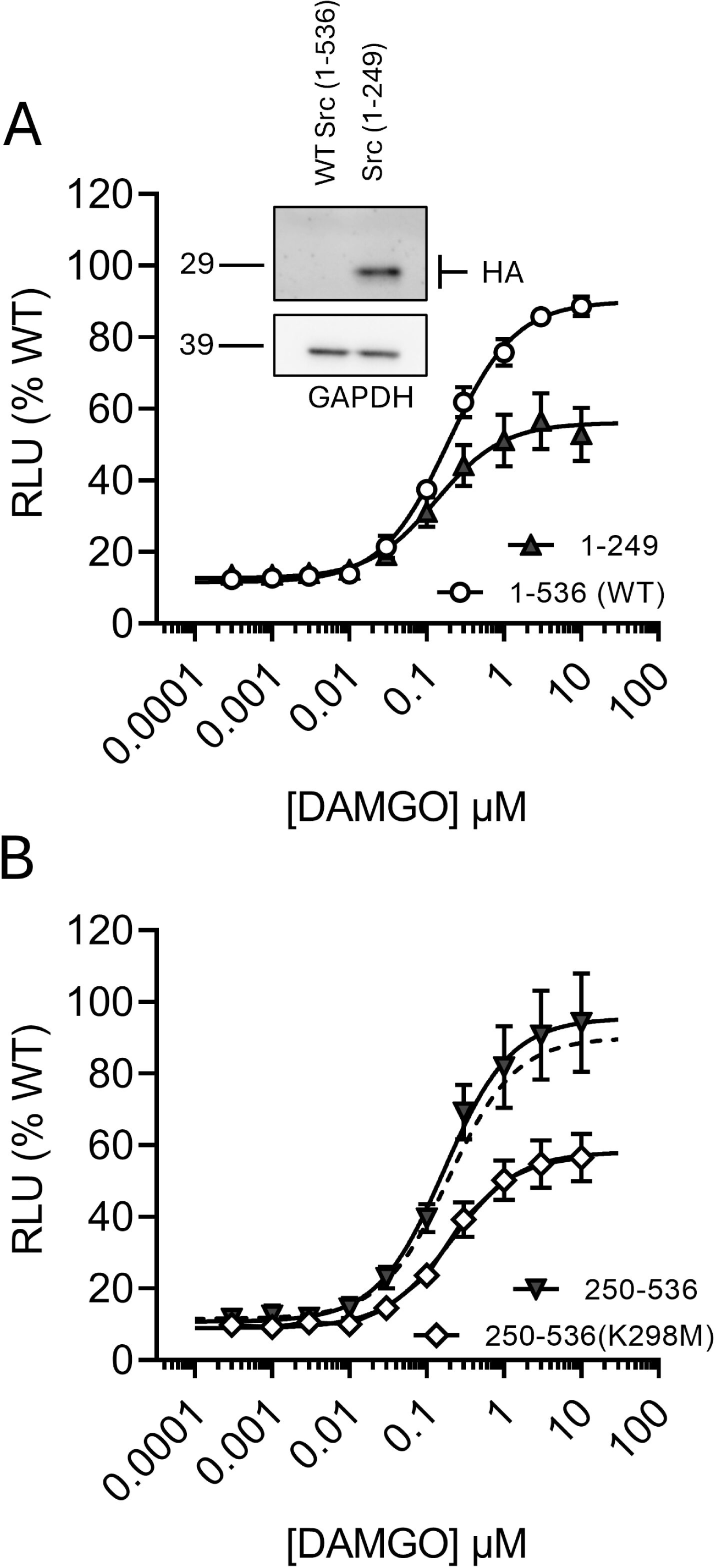
Transient overexpression of c-Src mutants lacking catalytic function reduce β-arrestin2 recruitment to μ receptors. Concentration-response relationships of DAMGO as a recruiter of β-arrestin2 to μ receptors in PathHunter CHO cells transiently overexpressing (**A**) full-length (residues 1-536; WT) versus C-terminally truncated (residues 1-249) human c-Src and (**B**) functional (residues 250-536) versus catalytically non-functional (residues 250-536(K298M)) N-terminally truncated human c-Src. The insert in (A) demonstrates overexpression of C-terminally truncated c-Src using an antibody against HA (1:1000). Overexpression of other c-Src mutants is depicted in Supplementary Figure 5A. WT responses in (B) are depicted as a dashed line. Data are expressed as a percentage of maximum luminescence produced by DAMGO in cells overexpressing WT c-Src (% WT). Data from individual replicates were plotted and fitted with logistics functions to derive efficacy (E_MAX_) and potency (EC_50_) parameters, presented in Table 2. Data in A and B are the mean ± SEM of 9 replicates.

**Table 2:**
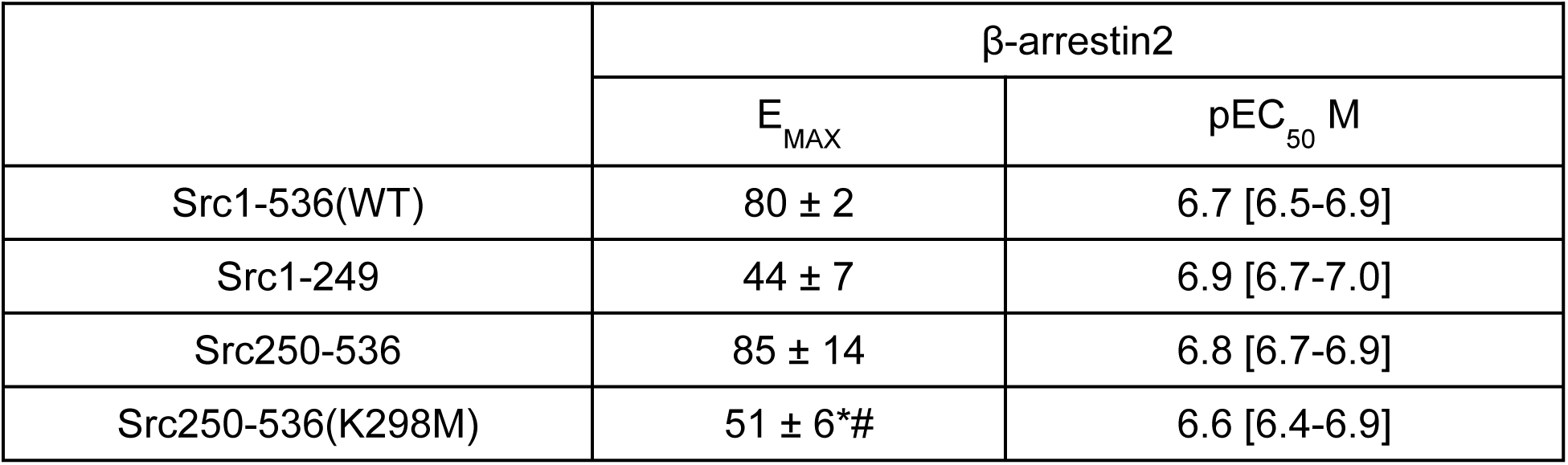
Efficacies and potencies of DAMGO to recruit β-arrestin2 to μ receptors in PathHunter CHO cells overexpressing wild type vs truncated mutant c-Src. Individual concentration-response relationships of DAMGO to recruit β-arrestin2 to μ receptors in cells overexpressing full length wild type (Src1-536), C-terminus truncated (Src1-249), N-terminus truncated (Src250-536) or N-terminus truncated kinase dead (Src250-536(K298M)) c-Src were fitted with a logistics function to yield efficacy and potency values (Figure 5A-B). A one-way ANOVA with posthoc Bonferroni correction revealed that the E_MAX_ of DAMGO to recruit β-arrestin2 was reduced by overexpression of Src1-249 and Src250-536(K298M) when compared to wild type c-Src (F3,30 = 6.4). Additional comparisons revealed a significant reduction in the E_MAX_ of DAMGO in cells overexpressing the kinase dead Src250-536KD when compared to functional kinase, Src250-536. By contrast, there was no impact of overexpressing any c-Src mutant on the potency of DAMGO (F3,30 = 2.8). *p < 0.05 when compared to wild type Src1-536 and # p < 0.05 when compared to Src250-536.

| | β-arrestin2 | |
| --- | --- | --- |
| | $E_{MAX}$ | $pEC_{50}$ M |
| Src1-536(WT) | $80 \pm 2$ | 6.7 [6.5-6.9] |
| Src1-249 | $44 \pm 7$ | 6.9 [6.7-7.0] |
| Src250-536 | $85 \pm 14$ | 6.8 [6.7-6.9] |
| Src250-536(K298M) | $51 \pm 6^*\#$ | 6.6 [6.4-6.9] |

Compared to cells overexpressing WT c-Src, overexpression of the N-terminal truncated mutant, Src1-249, reduced the E_MAX_ of DAMGO to recruit β-arrestin2 to μ receptors by half (Figure 5A), whereas DAMGO caused similar levels of β-arrestin2 recruitment in cells overexpressing the C-terminal truncated mutant, Src250-536 (Figure 5B; Table 2). We also exposed cells overnight to a phosphopeptide mimetic of the SH2 domain, caffeic acid-pYEEIE, that inhibits c-Src scaffolding in a competitive binding assay (Park et al., 2002). Caffeic acid-pYEEIE (200 nM) had no effect on the E_MAX_ of pEC_50_ of DAMGO or endomorphin-2 to recruit β-arrestin2 (Supplementary Figure 5B-C) confirming that non-catalytic functions of c-Src do not contribute to β-arrestin2 recruitment to μ receptors.

Focusing on the contribution of c-Src catalytic function, we also overexpressed a C-terminus mutant rendered catalytically inactive by methionine substitution of lysine at residue 298 (Src250-536(K298M)) to assess whether this mutant, which is dominant negative for internalisation of the β2-adrenergic receptor (Miller et al., 2000), diminishes β-arrestin2 recruitment in cells overexpressing C-terminus truncated Src. Indeed, Src250-536(K298M) reduced the E_MAX_ of DAMGO to recruit β-arrestin2 to μ receptors when compared to the functional Src250-536 mutant (Figure 5B; Table 2). Importantly, there was no change in viability between cells overexpressing WT Src compared to Src1-249, or between cells overexpressing Src250-536(K298M) compared to Src250-536 (Supplementary Figure 5D-E) demonstrating that the reduction in β-arrestin2 recruitment is not caused by lower cell densities. Taken together, these data demonstrate that the catalytic domain of c-Src participates in the agonist-evoked recruitment of β-arrestin2 to μ receptors.

We further tested the hypothesis that c-Src catalytic function participates in β-arrestin2 recruitment by overexpressing a c-Src catalytic mutant rendered constitutively active by phenylalanine substitution of tyrosine at residue 530 (SrcY530F) and assessing the efficacy and potency of DAMGO to recruit β-arrestin2 relative to cells overexpressing WT Src (Supplementary Figure 5F). DAMGO was significantly more efficacious as a recruiter of β-arrestin2 to μ receptors in cells overexpressing SrcY530F (111 ± 21%) compared to cells overexpressing WT (1-536) c-Src, although its potency was not affected (Supplementary Figure 5F). Furthermore, constitutive recruitment of β-arrestin2 in the absence of DAMGO was also enhanced by overexpression of SrcY530F to 162 ± 22% of the basal recruitment by WT c-Src (Supplementary Figure 5G), revealing that c-Src catalytic activity may also participate in constitutive, ligand-independent, recruitment of β-arrestin2.

### Overexpression of mutant c-Src constructs with no kinase activity abolishes the impact of c-Src inhibitors on β-arrestin2 recruitment to μ receptors

Since overexpression of c-Src mutants lacking kinase function diminishes β-arrestin2 recruitment to μ receptors (Figure 5A-B), we hypothesised that these mutants would also eliminate the impact of eCF506 and PP2 on DAMGO evoked recruitment of β-arrestin2 (Figure 3A). PathHunter CHO cells transiently overexpressing WT c-Src, or c-Src mutants lacking kinase activity (Src1-249 and Src250-536(K298M)), were exposed to eCF506, PP2 or PP3 overnight and compared to cells overexpressing the same construct but exposed to an equal volume of DMSO (i.e., the maximum recruitment in these cells). Mean efficacy (E_MAX_) and potency (pEC_50_) values of DAMGO to recruit β-arrestin2 are summarised in Supplementary Table 3.

Exposure to PP2 or eCF506 (300 nM) reduced the E_MAX_ of DAMGO to recruit β-arrestin2 to μ receptors in PathHunter CHO cells transiently overexpressing WT c-Src (F3,16 = 19.6), whereas PP3 (300 nM) had no effect (Figure 6A). However, only eCF506 limited the E_MAX_ of DAMGO in cells transiently overexpressing c-Src1-249 (F3,16 = 3.4; Figure 6B), although its inhibition was diminished (Supplementary Table 3), and neither eCF506 nor PP2 caused a significant reduction in the E_MAX_ of DAMGO in cells transiently overexpressing c-Src250-536(K298M) (F3,19 = 1.2; Figure 6C). There was no effect of c-Src inhibition on the potency of DAMGO in cells overexpressing WT c-Src, Src1-249 or Src250-536(K298M) (Supplementary Table 3). Therefore, these data demonstrate that the restriction of DAMGO’s E_MAX_ by PP2 and eCF506 is mediated through inhibition of c-Src kinase catalytic function.

**Figure 6:**
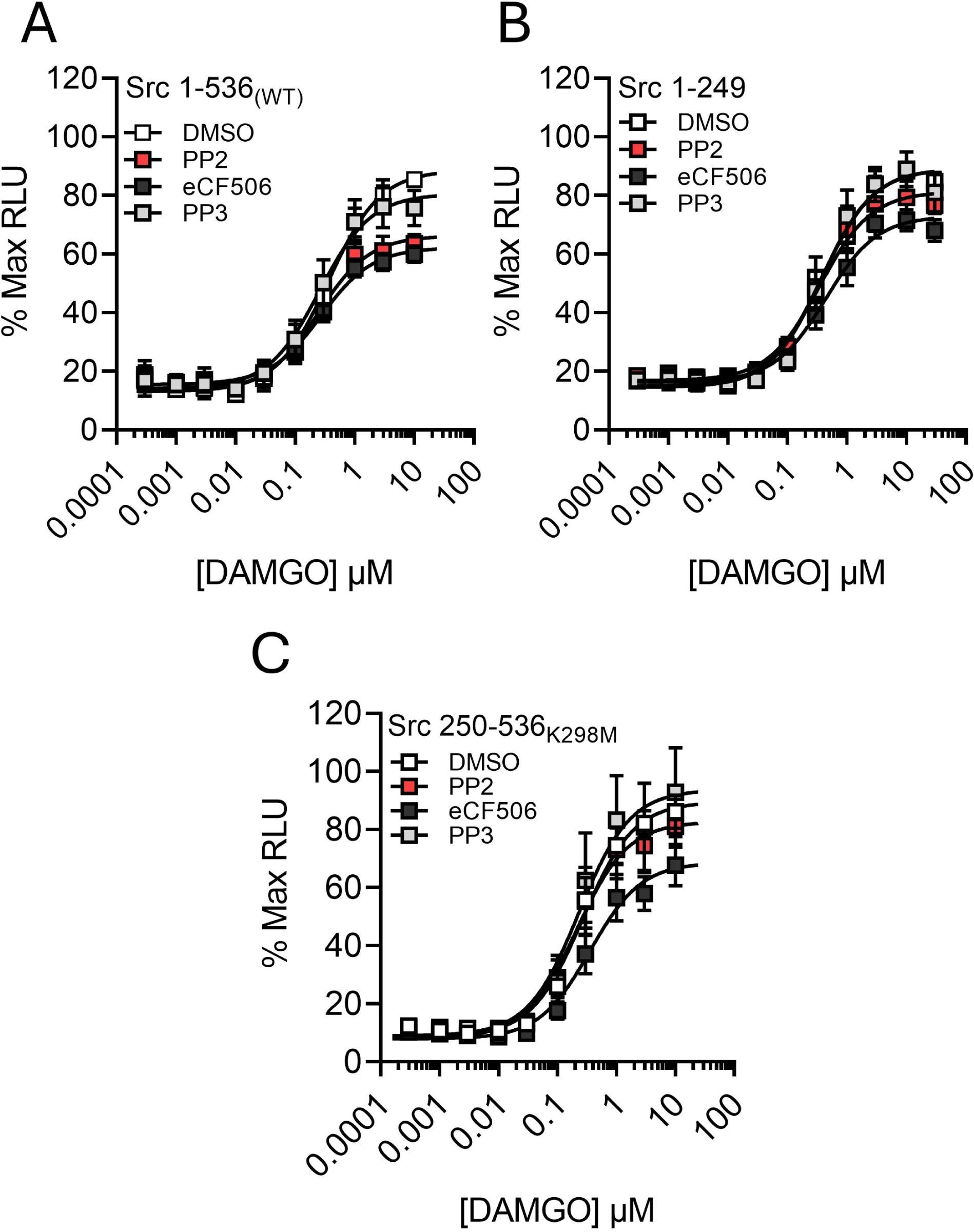
Catalytically inactive c-Src mutants abolish inhibitor effects on β-arrestin2 recruitment. Concentration-response relationships of DAMGO to recruit β-arrestin2 to human μ receptors in PathHunter CHO cells transiently overexpressing (**A**) full-length (residues 1-536; WT), (**B**) C-terminally truncated (residues 1-249) or (**C**) N-terminally truncated and catalytically inactive (residues 250-536(K298M)) human c-Src following 16-h exposure to PP2, PP3, eCF506 (300 nM) or an equal volume of DMSO. Data are expressed as a percentage of maximum luminescence (% Max RLU) produced by DAMGO in DMSO exposed cells present in duplicate on each plate i.e., cells overexpressing (A) WT, (B) Src 1-249 or (C) Src 250-536(K298M) are each expressed relative to the maximum β-arrestin2 recruitment of each construct caused by DAMGO. Data from individual replicates were plotted and fitted with logistics functions to derive efficacy (E_MAX_) and potency (EC_50_) parameters, presented in Supplementary Table 3. Data are the mean ± SEM of 5-6 replicates.

### c-Src inhibition by eCF506 enhances μ receptor surface availability and GRK2 expression

Activated μ receptors are targeted for c-terminal phosphorylation by GRKs, which enhances the affinity for β-arrestin2 binding (Williams et al., 2013). Therefore, a possible explanation for the reduction in β-arrestin2 recruitment to μ receptors caused by inhibition of c-Src is a downregulation in the expression of μ receptors, β-arrestin2 and/or GRKs. We investigated this using HEK cells stably expressing recombinant human μ receptors tagged with HiBiT (Supplementary Figure 6A). Importantly, this tag did not disrupt μ receptor function when assessed using the cAMP accumulation assay (Supplementary Figure 6B).

Overnight exposure to eCF506 (1 µM) elevated surface luminescence by 16 ± 3% (range 7 – 24%, n = 6) of that in HiBiT-Oprm1 cells exposed to an equal volume of DMSO (Supplementary Figure 6C). Western blot of HiBiT-Oprm1 cell lysates following overnight exposure to eCF506 (1 nM – 1 µM) also revealed a concentration-dependent increase in the expression of GRK2 (Supplementary Figure 6D). Neither GRK5 expression nor β-arrestin2 expression were affected (Supplementary Figure 6E-F).

### c-Src inhibition by eCF506 has no effect on μ receptor c-terminal phosphorylation

Agonist-evoked phosphorylation of the μ receptor is both agonist and GRK-dependent with lower efficacy agonists primarily causing GRK5/6 mediated phosphorylation of μ receptors restricted to S375 while higher efficacy agonists recruit GRK2/3 to phosphorylate S375 and additional distal residues T370, T376 and T379 (Gillis et al., 2020; Underwood et al., 2024). Therefore, we assessed whether upregulation of GRK2 caused by eCF506 (Supplementary Figure 6D) altered c-terminal phosphorylation of μ receptors using well established phosphosite-specific antibodies against residues T370, S375, T376 and T379 (Gillis et al., 2020; Fritzwanker et al., 2021; Underwood et al., 2024).

DAMGO (1 nM - 10 μM) caused a concentration-dependent increase in the phosphorylation of residues T370, S375, T376 and T379 of HiBiT-tagged human μ receptors, significantly elevating phosphorylation after exposure to ≥100 nM when compared to cells without DAMGO (Figure 7A-D; Supplementary Figure 7). eCF506 (1 μM) had no impact on the level of phosphorylation caused by DAMGO demonstrating that c-Src inhibition does not affect agonist-evoked phosphorylation of the μ receptor c-terminus.

**Figure 7:**
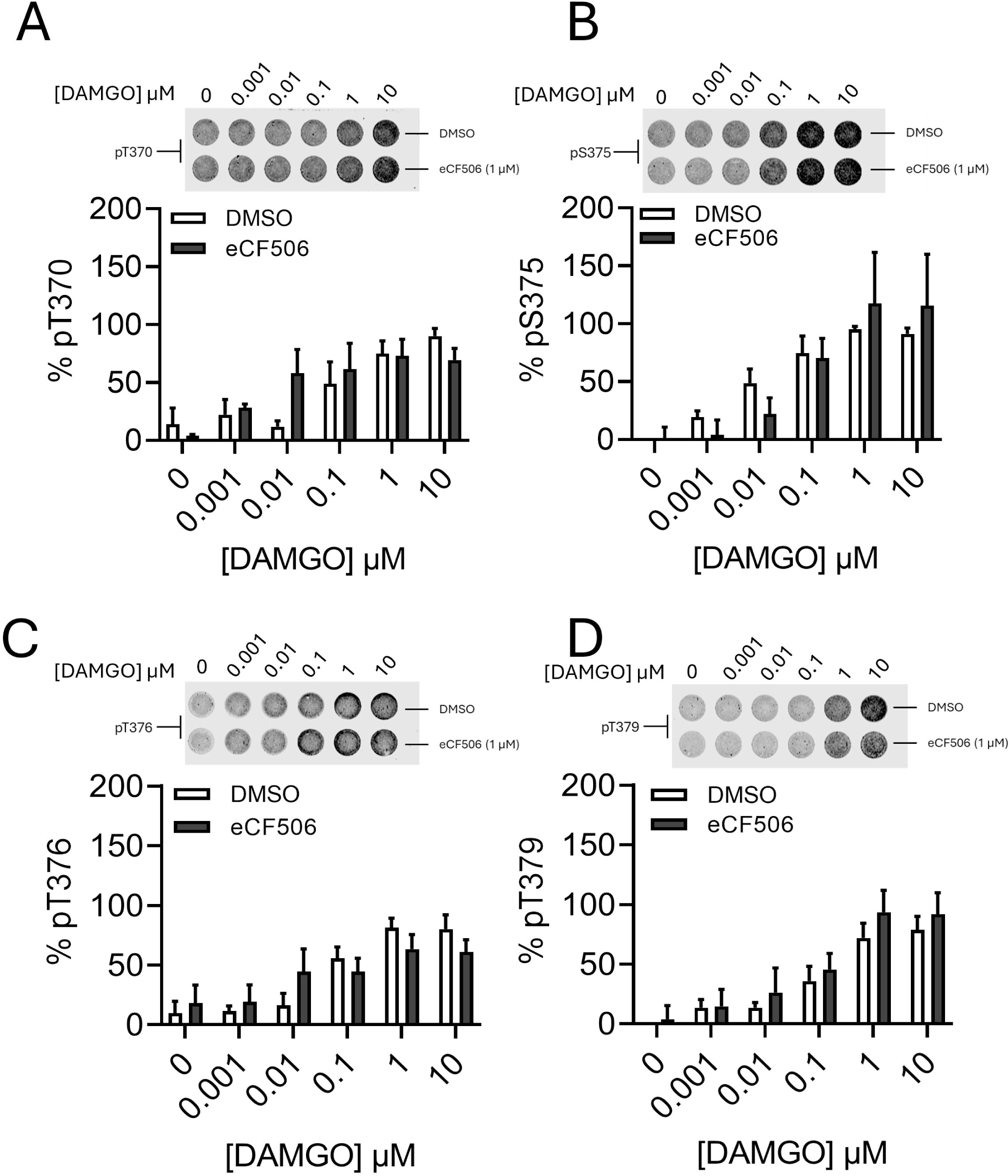
eCF506 does not influence μ receptor c-terminal phosphorylation. DAMGO evoked c-terminal phosphorylation at residues T370, S375, T376 and T379 of HiBiT-tagged μ receptors expressed in HEK cells was assessed using an in-cell western blot approach. The inserts show representative fluorescence in the phosphorylated μ receptor channel (700 nm). Fluorescence intensities in the non-phosphorylated μ receptor channel, determined by a monoclonal antibody against the HiBiT tag, are presented in Supplementary Figure 7. Data are expressed as a ratio of phosphorylated residue to total μ receptor in each well and subsequently as a percentage of maximum phosphorylation caused by DAMGO in HEK cells exposed to DMSO. (**A-D**) Fluorescent analysis of DAMGO (15 minutes at 37 °C) evoked phosphorylation at (**A**) T370, (**B**) S375, (**C**) T376 and (**D**) T379 in HiBiT-μ receptor expressing HEK cells exposed to 16 h eCF506 (1 μM) or an equal volume of DMSO. A two-way ANOVA was used to compare treatment (DMSO vs eCF506) and DAMGO concentration for each phosphorylated residue. There was a significant effect of DAMGO concentration on the phosphorylation of (**A**) pT370 (F5,35 = 8.5), (**B**) pS375 (F5,42 = 7.9), (**C**) pT376 (F5,36 = 9.4) and (**D**) pT379 (F5,48 = 14.2) although there was no effect of eCF506 on the phosphorylation of any residue. Pairwise comparisons with the Dunnett posthoc correction applied revealed that μ receptor phosphorylation at residues T370, S375, T376 and T379 were significantly elevated at concentrations ≥ 100 nM DAMGO when compared to control (no DAMGO). See Methods for details of in-cell western blotting conditions and visualisation. Data are the mean ± SEM of 4 replicates.

**Figure 8:**
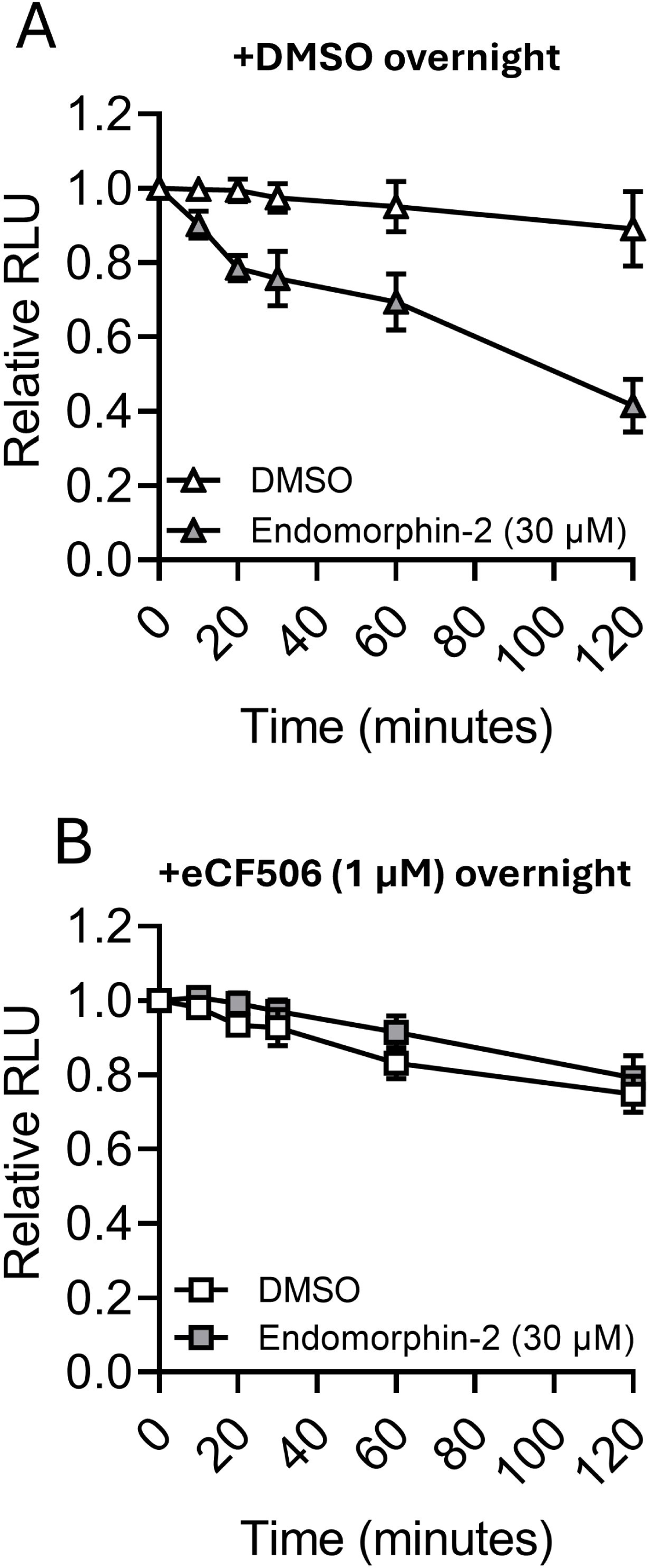
eCF506 abolishes endomorphin-2 evoked μ receptor internalisation. HEK293 cells stably expressing HiBiT-tagged human μ receptors were used to assess the impact of c-Src inhibition by eCF506 on endomorphin-2 (30 μM) induced internalisation. (**A**) Endomorphin-2 caused a reduction in luminescence corresponding to a reduction in cell-surface μ receptor availability across 2-hours in cells exposed to DMSO overnight when compared to the luminescence produced by cells exposed to the vehicle, DMSO. (**B**) By contrast, endomorphin-2 failed to cause a reduction in cell-surface μ receptor availability in cells exposed to eCF506 (1 μM). Data are the mean ± SEM of n = 4 replicates and expressed relative to the luminescence recorded immediately after agonist dilution.

### c-Src inhibition by eCF506 abolishes endomorphin-2 evoked internalisation of μ receptors

c-Src participates in β2-adrenergic receptor internalisation by targeting arrestin bound receptors to clathrin-mediated endocytosis machinery (Luttrell et al., 1999). Therefore, we assessed whether c-Src inhibition by eCF506 similarly influences the internalisation of μ receptors using a cell-surface luminescence assay in HEK cells stably expressing HiBiT-tagged human μ receptors. Our reference agonist, DAMGO, was revealed to complement with LgBiT (Supplementary Figure 8). Therefore, cells were exposed to eCF506 (1 μM) or an equal volume of DMSO overnight and the surface availability of μ receptors was assessed in real time in the presence of endomorphin-2 (Figure 8).

Endomorphin-2 (30 μM) caused a gradual reduction in μ-opioid receptor surface expression over 2 hours, reaching 42 ± 7% of the baseline luminescence in cells pretreated overnight with DMSO. In contrast, receptor surface levels remained largely unchanged (89 ± 10%) in vehicle-treated cells (Figure 8A). By contrast, endomorphin-2 failed to cause μ receptor internalisation in cells exposed to eCF506 (Figure 8B). The luminescence after 2-hours remained at 75 ± 5% and 79 ± 6% in eCF506 treated cells treated with vehicle or endomorphin-2, respectively. These data demonstrate that c-Src activation is required for internalisation of μ receptors by endomorphin-2.

## Discussion

Consistent with the properties of less selective Src inhibitors, the type 1.5 conformationally selective inhibitor, eCF506, dose-dependently attenuated morphine antinociceptive tolerance in mice and caused a transient recovery from established tolerance. Furthermore, in cellular models, eCF506, PP2 and targeted degradation of c-Src diminished the recruitment of β-arrestin2 to agonist-activated μ receptors, whereas PP3, a c-Src inactive analogue of PP2, had no effect. β-arrestin2 recruitment was also reduced in cells overexpressing catalytically inactive c-Src mutants (Miller et al., 2000), and neither PP2 nor eCF506 limited the efficacy of DAMGO in cells overexpressing a dominant negative kinase (K298M). By contrast, overexpression of a c-Src Y530F mutant construct with enhanced catalytic function (Roskoski, 2005) increased β-arrestin2 recruitment to μ receptors by DAMGO and led to constitutive recruitment.

The impact of c-Src inhibition on limiting β-arrestin2 recruitment also coincided with a failure of endomorphin-2 to cause endocytosis and elevated surface availability of μ receptors. The latter is consistent with prior observations in mouse DRG neurones exposed to PP2 (Walwyn et al., 2007), demonstrating an indispensable role of c-Src in μ receptor internalisation similar to its regulation of other GPCRs (Miller et al., 2000; Hong et al., 2009).

β-arrestin2 is likely required to target c-Src to activated μ receptors as evidenced by disrupted c-Src distribution and an inability of DAMGO to phosphorylate c-Src in the DRG neurones of β-arrestin2-/- mice (Walwyn et al., 2007). In keeping with this sequence of events, exposure to μ receptor agonists caused phosphorylation of c-Src that correlated with agonist efficacy for recruiting β-arrestin2. These findings are consistent with the idea that arrestins recruit c-Src via its SH3 domain following engagement with the phosphorylated receptor tail (Luttrell et al., 1999; Yang et al., 2018; Pakharukova et al., 2021). However, our finding that pharmacological inhibition, targeted c-Src degradation, and overexpression of non-functional mutant constructs of c-Src also limit β-arrestin2 recruitment to μ receptors, suggests additional roles of c-Src in regulating μ receptor signal transduction upstream of β-arrestin2 binding. Indeed, c-Src can directly phosphorylate μ receptors (Zhang et al., 2017) and this occurs in mouse embryonic fibroblasts depleted of arrestins (Zhang et al., 2009).

Early evidence reveals that c-Src-mediated phosphorylation of GRK2 enhances its catalytic activity (Sarnago et al., 1999), whilst interactions between c-Src and arrestins can promote GRK2 degradation (Penela et al., 2001). It is therefore consistent that c-Src inactivation through pharmacological inhibition by eCF506 resulted in elevated GRK2 expression. However, it is interesting that upregulation of GRK2 by eCF506 had no impact on the phosphorylation barcode evoked by DAMGO at μ receptors; several serine/threonine residues present on its c-terminus are well established to be specific targets of GRK2/3 (Just et al., 2013; Miess et al., 2018; Drube et al., 2022; Underwood et al., 2024). This includes T376 and T379 which are not phosphorylated by morphine (Gillis et al., 2020; Just et al., 2013). It is possible that high basal expression of GRK2/3 in HEK cells (Atwood et al., 2011; Just et al., 2013) is sufficient to saturate phosphorylation of μ receptors in our system when stimulated with high efficacy agonists like DAMGO. However, it is important to note that c-terminal phosphorylation of μ receptors does not always correlate with agonist efficacy. Some high efficacy agonists like SR-17018 cause substantial GRK2/3 mediated phosphorylation but have limited efficacy for β-arrestin2 recruitment (Fritzwanker et al., 2021; Gillis et al., 2020; Schmid et al., 2017; Singleton et al., 2024). In this context, μ receptors adopt a conformation less permissive to β-arrestin2 signalling (Singleton et al., 2024; Stahl et al., 2021).

Regardless of the order of events, inhibition of c-Src catalytic function impairs β-arrestin2 recruitment to μ receptors and this may have important implications for improving the long-term use of opioid analgesics. Compelling evidence has confirmed that β-arrestin2 recruitment to μ receptors is responsible for the development of analgesic tolerance to morphine (Bohn et al., 2000; Yang et al., 2011; Bull et al., 2017; Kliewer et al., 2019). Our finding that c-Src catalytic activity is required for β-arrestin2 recruitment provides mechanistic insights into how eCF506 diminishes morphine analgesic tolerance. Our prior study also reveals that the c-Src/c-Abl kinase inhibitor, dasatinib, and PP2 abolish tolerance to morphine (Bull et al., 2017). These bind c-Src in its active conformation (Tokarski et al., 2006; Muratore et al., 2009) consistent with a requirement of c-Src catalytic function in the development of tolerance. It is unclear whether these effects are solely mediated through restricted β-arrestin2 recruitment, although several phenotypes in DRG neurones from β-arrestin2-/- mice, including elevated surface expression of μ receptors and their constitutive coupling to VACCs, are mimicked in wild type DRGs treated with c-Src inhibitors (Walwyn et al., 2007). Nevertheless, some behaviours caused by the absence of β-arrestin2, including prolongation of basal tail latencies and enhanced potency of morphine to cause antinociception (Bohn et al., 2000; Lam et al., 2011) are not observed in wild type mice treated with eCF506. This implies that disruption to c-Src signalling alone is insufficient to recapitulate these phenomena. Interestingly, eCF506 had only transient effects on restoring morphine analgesia in μ+/- mice rendered tolerant by repeated exposure to morphine. This may be explained by a temporary increase to the surface availability of μ receptors revealed in the HiBiT assay. While eCF506 abolishes radiation-induced lung fibrosis, it is ineffective at reversing established lung pathologies (Choi et al, 2025). Therefore, prophylactic targeting of c-Src by eCF506 may be required for optimising its efficacy *in vivo*.

Beyond its role in the development of analgesic tolerance to morphine, mounting evidence supports a broader detriment of c-Src activity on pain and analgesia. c-Src is phosphorylated and co-precipitates with μ receptors during the remission phase of inflammatory hypersensitivity (Walwyn et al., 2016) and c-Src inhibition protects against reinstatement of mechanical hypersensitivity caused by the μ receptor inverse agonist naltrexone (Chen & Marvizón, 2020). Several groups also report a protective effect of dasatinib and PP2 on inflammatory, neuropathic and cancer-induced pain outcomes (Liu et al., 2008; Appel et al., 2017; Ge et al., 2020; Singleton and Hales, 2023). Our prior study demonstrated that dasatinib abolishes the development of morphine-induced mechanical hypersensitivity (Singleton and Hales, 2023).

Taken together, our findings demonstrate an important contribution of c-Src catalytic function in the recruitment of β-arrestin2 to μ receptors and provide a mechanism by which c-Src inhibitors may confer resistance against the development of antinociceptive tolerance to morphine. We establish that inhibition of the c-Src catalytic function increases the surface availability of μ receptors whilst diminishing β-arrestin2 recruitment and μ receptor internalisation without affecting G-protein coupling. These properties mimic the benefits sought by the development of biased agonists, which tend to be partial agonists, while maintaining full opioid efficacy. Inhibition of c-Src may provide a unifying strategy to limit hypersensitivity and improve the long-term analgesic effects of opioids that can be leveraged to better treat chronic pain.

## Methods

### Animals

Male and female wild type (WT) and μ+/- C57BL/6 mice, weighing 24 – 30 g and aged 10 – 14 weeks at the start of each experiment, were bred in house in a temperature (19-24°C) controlled unit on a 12-h light cycle at the University of Dundee. Free access to food and water was available throughout all studies. Behavioural experiments took place in a separate room that mice were habituated to for 3 days during handling prior to starting each experiment. Genotypes were confirmed at the end of the experiment using the automated genotyping service provided by Transnetyx (Cordova, USA). Mice were housed in social groups of 3 or more same-sex littermates in open-top opaque cages enriched with polycarbonate tubes and cable ties and provided with routine nesting and bedding material. All procedures were carried out under the authority of licences granted by the UK Home Office under the Animals (Scientific Procedures) Act 1986 and approved by the University Welfare and Ethical Use of Animals Committee, acting in its capacity as an Animal Welfare and Ethical Review Body as required under the Act. Animal studies are reported in compliance with the ARRIVE guidelines 2.0 (Percie du Sert et al., 2020).

### Tail withdrawal assay

Morphine antinociception and the development of antinociceptive tolerance was measured using a warm water tail withdrawal assay. This is a well-established assay that measures analgesia and the development of tolerance through spinal reflexes in rodents (Deuis et al., 2017). Mice were restrained in polycarbonate tubes present in home cages and the distal third of their tails submerged into warm (48 ± 0.1°C) water (thermostatic circulator bath Optima general purpose 12 L stainless steel tank; Fisher Scientific, UK). Tail withdrawal latencies were measured up to a maximum of 15 seconds per assessment to avoid tissue injury.

Withdrawal latencies were measured before and 30 minutes after administration (p.o.) of eCF506 (20 mg/kg or 80 mg/kg) or vehicle (ultrapure water with 5% DMSO and 5% Cremophor). Morphine (up to 30 mg/kg) was injected (s.c.) immediately after the second latency assessment and tail withdrawal latencies reassessed 30 minutes later. For dose-response relationships on day 1 and day 10, all mice received escalating doses of morphine (0.1 – 30 mg/kg) with each injection separated by 30 minutes. Tail withdrawal latencies were reassessed 30 minutes after each injection using the same approach.

Morphine antinociception is presented as a percentage of maximum possible effect, determined using the equation:

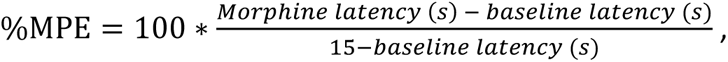

Where baseline latency represents the time to withdraw (s) in the absence of eCF506 or matched vehicle treatment. ED_50_ values for each animal were derived from a logistic function fitted to individual dose–response relationships and constrained to a maximum %MPE of ≤100%.

### Sample size estimate

Data from a prior study (Bull et al., 2017) assessing the impact of c-Src inhibition by dasatinib (5 mg/kg; i.p.) were used to establish appropriate sample sizes. Power analysis revealed an effect size (d) of 2.3. Our prior studies (Bull et al., 2017; Singleton et al., 2023) revealed no influence of sex in the tail withdrawal assay. Therefore, analyses were performed without sex as an experimental variable and groups contained an even distribution of males and females. In experiments performed in μ+/- mice, entire litters from 2 heterozygous breeding pairs were included to avoid surplus breeding whilst achieving adequate power. One μ+/- mouse was removed during the study (barbering), resulting in a sample size of n = 7 (vehicle: 3M + 4F) and n = 8 (eCF506: 3M + 5F). Mice were randomly allocated to receive either vehicle or eCF506 prior to the start of each experiment and tail withdrawal latencies were assessed by a second individual blinded to treatment group. Data were unblinded following statistical analysis of dose-response relationships to establish morphine

potency (ED_50_) and the development of tolerance. All mice completing each study were included in data analysis.

## Materials

eCF506 was synthesised in house at the University of Edinburgh as previously described (Fraser et al., 2016; Temps et al., 2021). DAMGO ([D-Ala2, N-MePhe4, Gly-ol]-enkephalin) and forskolin (HelloBio, UK), morphine sulfate salt pentahydrate (#M8777), fentanyl citrate (#F3886), oxycodone hydrochloride (#BP1068; all from Merck, UK), TRV130 (#MBS3601671; CliniSciences, UK), buprenorphine hydrochloride (#2808), β-FNA (#0926), PP2 (#1407), PP3 (#2794), caffeic acid pYEEIE (#1935; all from Tocris, UK), DAS-5-oCRBN (#HY-163144; Cambridge Biosciences, UK) were prepared as 10 mM (DAMGO, morphine, fentanyl, buprenorphine, TRV130, PP2, PP3, DAS-5-oCRBN, caffeic acid-pYEEIE and eCF506), 20 mM (oxycodone), 25 mM (forskolin) or 100 mM (β-FNA) stock solutions in DMSO (DAMGO, forskolin, buprenorphine, β-FNA, PP2, PP3, DAS-5-oCRBN, caffeic acid-pYEEIE and eCF506) or water (fentanyl, morphine, oxycodone). Morphine for *in vivo* administration was prepared as a 2 mg/ml solution in 0.9% normal saline. eCF506 was prepared as a 10 mg/ml solution in warm ultrapure water and supplemented with 2% Kolliphor EL. Matched vehicle administrations contained the same constituents without eCF506. All compounds administered to mice were prepared in an aseptic environment and 0.22 μm filter sterilised (#SCGP00525; Merck, UK) immediately before use on each day.

### Cell culture

PathHunter CHO-K1 (RRID:CVCL_KY70) and U2OS (RRID:CVCL_LB00) OPRM1-ARRB2 cells stably overexpressing β-galactosidase enzyme tagged human μ receptors and β-arrestin2 (DiscoverX, US) and Human Embryonic Kidney-293 (HEK293; RRID:CVCL_0045) cells were used throughout this study. HEK293 and PathHunter CHO-K1 cells were maintained in DMEM (the latter with F12) while PathHunter U2OS cells were maintained in McCoys 5A medium. All media were supplemented with GlutaMAX™, foetal bovine serum (10% v/v), penicillin (5000 U/ml) and streptomycin (5000 μg/ml). PathHunter cells were additionally supplemented with selection agents geneticin (500 μg/ml) and hygromycin B (250 μg/ml) to maintain stable expression of μ receptors and β-arrestin2. A stably transfected HEK293-HiBiT-Oprm1 cell line was generated using a 10-day exposure to geneticin (800 μg/ml) and limiting dilution followed by single-cell expansion and maintained with geneticin (500 μg/ml). All cells were grown in a humidified incubator at 37 °C in 5% CO_2_ and routinely sub-cultured when they reached 80% confluence.

### Transfection

Cells were plated for transfection in 35 mm Nunc™ dishes (#150318; ThermoFisher Scientific, UK) in 2 ml Opti-MEM at a density of 1x10^6^ cells per dish and incubated overnight at 37°C. The following day, 2 μg cDNA encoding c-Src mutant constructs or the cAMP sensitive plasmid, pGloSensor-22F, were transfected in 200 μl of Opti-MEM using lipofectamine-2000 (#11668019; ThermoFisher Scientific, UK) at a ratio of 2.5:1 (lipofectamine:cDNA) and returned to the incubator overnight. Cells transiently overexpressing c-Src proteins or pGloSensor were harvested for functional assays the next day. Parallel transfections using 2 μg of eGFP were performed under identical conditions and visualised using fluorescence microscopy to verify successful transfection and monitor transfection efficiencies.

### β-arrestin2 recruitment

β-arrestin2 recruitment was assessed using a protein complementation assay (Eurofins DiscoverX, UK) in PathHunter CHO or U2OS cells stably overexpressing human μ receptors and β-arrestin2 each fused to complementary fragments of β-galactosidase. Cells were plated at a density of 5000 cells per well in 50 μl of Opti-MEM in 96-well half area white bottom plates (Greiner #675083) and left to settle overnight. Opti-MEM was replaced the following morning with agonists prepared in fresh Opti-MEM then incubated at 37°C for 90 minutes. DiscoverX chemiluminescence substrates (Galacton Star and Emerald-II) were reconstituted as recommended in Opti-MEM and added to each well then incubated for 2 h at room temperature in the dark. Luminescence was measured on a GloMAX Navigator luminometer (Promega, UK) using a 1-s integration time. Data were measured in technical duplicates on each plate and the average of these replicates recorded as n = 1.

### cAMP accumulation

Cells transiently expressing the pGloSensor protein were plated in the same 96-well half area white bottom assay plates at a density of 15000 cells per well in 50 μl of Opti-MEM and left to settle overnight. The following morning, Opti-MEM was replaced with 40 μl of HEPES buffered HBSS containing luciferin (3 mM) and forskolin (30 μM) at pH 7.4 and incubated in the dark at room temperature for 2 hours. Luminescence was measured on a GloMAX Navigator luminometer using a 1-s integration time prior to the addition of 10 μl μ receptor agonists (made up at 5x concentration in the same luciferin and forskolin containing buffer) into each well. Luminescence was reassessed 30 minutes after agonist dilution using the same approach. Data were measured in technical duplicates on each plate and the average of these replicates recorded as n = 1.

For experiments involving a reduction in μ receptor availability, the number of available μ receptors was restricted using the irreversible μ receptor antagonist, β-funaltrexamine (β-FNA). Cells were exposed to β-FNA (100 nM) for 1 hour at 37°C as our prior studies have consistently revealed that this concentration is sufficient to abolish μ receptor reserve of DAMGO in the cAMP accumulation assay (Singleton et al., 2021; Singleton et al., 2024). β-FNA was reconstituted in luciferin and forskolin containing buffer and exposed to cells for the first hour of the 2-hour incubation. Cells were washed to remove unbound β-FNA and replenished with 40 μl of fresh assay buffer before assessing luminescence 1 hour later. Agonist addition and subsequent measurements were performed as described above.

### Inhibition of c-Src

For experiments involving additional overnight incubation with c-Src inhibitors/degraders or vehicle controls, these were prepared at 2x concentration in Opti-MEM and 50 μl added directly to each well at the end of the day before each assay. β-arrestin2 recruitment and inhibition of cAMP accumulation were assessed the following day using the same approaches described above with the exception that c-Src modulators were additionally included in the agonist (β-arrestin2 assay) or luciferin and forskolin (cAMP assay) incubation steps.

### Cell viability

Cell viability was measured using an MTT or CellTiter-Glo assay (#G7570; Promega, UK) at the start of each experiment. In both assays, cells were seeded in Opti-MEM and treated as described in the overnight c-Src exposure paradigm then left to settle overnight. The following morning, when measuring MTT absorbance, Opti-MEM was replaced with 100 μl MTT (0.5 mg/ml) reconstituted in dPBS and incubated at 37°C for 3 hours. MTT was then replaced with 100 μl isopropanol containing HCl (4 mM) and 0.1% (v/v) NP40 and incubated in the dark with agitation for 15 minutes. Absorbance was measured at 590 nm using a FlexStation-III multimodal plate reader (Molecular Devices, USA). In the CellTiter-Glo assay, a ladder of known cell densities (625-20000 per well) without exposure to treatment was also included on each 96-well half area white plate to extrapolate cell plating efficiency. The following morning, Opti-MEM was replaced with 50 μl of fresh Opti-MEM followed by 50 μl of proprietary CellTiter-Glo substrate reconstituted as recommended. Plates were incubated at room temperature in the dark for 10 minutes and luminescence measured on a GloMAX Navigator luminometer using a 1-s integration time.

### Receptor tagging

Human ORF HaloTag® Oprm1 in pFN21A (clone #FHC11924) was digested with AsiSI and PmeI and the purified Oprm1 insert fragment was ligated into the N-terminus HiBiT tagged pFN39K vector

(Promega, UK) digested with AsiSI and PmeI. The final plasmid contained the IL-6 signal sequence (MNSFSTSAFGPVAFSLGLLLVLPAADPAP) for membrane insertion, followed by HiBiT (VSGWRLFKKIS) and a double GSSG-linker at the N-terminus of human Oprm1. The full open reading frame of the fusion protein was sequenced at the University of Dundee MRCPPU.

### Surface expression and internalisation assay

Cell surface availability and internalisation of human μ receptors were assessed using the HiBiT extracellular detection system (Promega, UK). Luminescence in this assay corresponds to the surface receptor density enabling real time measurements of receptor internalisation (Boursier et al., 2020). HEK cells stably expressing HiBiT-tagged human μ receptors were seeded in 96-well half area white plates (Greiner #675083) at a density of 10000 cells per well in Opti-MEM (50 μl) and exposed to eCF506 (1 μM reconstituted at 2x concentration in Opti-MEM) or an equal volume of DMSO overnight whilst incubated at 37°C and 5% CO_2_. The following morning, Opti-MEM was replenished with 40 μl of HiBiT extracellular detection buffer containing LgBiT (1:200) and furimazine (1:100), with either DMSO or eCF506 (1 μM). Plates were immediately placed into a GloMAX Navigator luminometer (Promega, UK) and luminescence assessed after 15 minutes using a 0.1-s integration time.

Internalisation of HiBiT-tagged human μ receptors was measured in the same experiment using the same plates. After baseline cell-surface luminescence was assessed, each well was diluted with 10 μl of endomorphin-2 (30 μM prepared at 5x concentration) or an equal volume of DMSO prepared in the same extracellular detection buffer. Plates were returned to the luminometer and luminescence assessed immediately and automatically at intervals across 2 hours. Data are expressed relative to the luminescence recorded immediately after agonist dilution. Data were measured in technical triplicates on each plate and the average of these replicates recorded as n = 1.

### Western blot

Cells were enzymatically detached using tryspin (0.25%) and pelleted by centrifugation at 3,200 x g for 2 minutes. Pellets were washed in dPBS and re-centrifuged to remove residual trypsin then incubated on ice for 30 minutes in RIPA buffer containing Roche cOmplete™ EDTA-free protease and phosphatase (PhosSTOP™) inhibitors with frequent agitation. Cell lysates were centrifuged for 10 minutes at 12,000 x g and protein concentration of the supernatant established using the Pierce BCA assay. Cell lysates (12 μg prepared in 12 μl of SDS reducing buffer with 20% β-mercaptoethanol) were boiled for 5 minutes at 95°C then separated by electrophoresis on NuPAGE™ 4-12% bis-tris polyacrylamide gels using MOPS SDS running buffer for 90-120 minutes at 120 mV. Proteins were transferred onto Cytiva Amersham™ 0.45-μm nitrocellulose membranes (Fisher scientific, UK) for 85 minutes at 200 mAh submerged in ice using tris-glycine transfer buffer supplemented with methanol (20% v/v) and SDS (3 mM). Membranes were blocked for 1 hour at room temperature in tris-buffered saline with 0.1% tween-20 (TBS-T) containing 5% (w/v) non-fat dried milk powder or 5% bovine serum albumin (BSA; Merck #A7906 >98%). Primary antibodies were diluted in the same blocking buffer and incubated overnight at 4°C. The following day, antibodies were replaced with compatible HRP-conjugated secondary antibodies diluted in the same blocking buffer for 1 hour at room temperature. Membranes were incubated with chemiluminescent substrate and visualised on a ChemiDoc multimodal imager.

### Antibodies

Polyclonal antibodies against c-Src (CST #2018S) or c-Src phosphorylated at residue Y416 (CST #2101) were purchased from Cell Signaling Technology (USA). Polyclonal antibodies against GAPDH (#ab9485 were purchased from Abcam (USA). Polyclonal antibodies against β-arrestin2 (#10171-1-AP) were purchased from Proteintech (U.K). Polyclonal antibodies against GRK2 (#AF4339) and against GRK5 (#AF4539) were purchased from R&D Systems (UK). Monoclonal antibodies against HiBiT (#N7200) were purchased from Promega (UK). Polyclonal antibodies raised against phosphorylated residues on μ receptors (T370, S375, T376, T379) were purchased from 7TM Antibodies (Jena, Germany; #7TM0319-SP). Primary antibodies were raised in rabbit (pSrc-Y416, Src, GAPDH, β-arrestin2 and phosphorylated μ receptor residues T370, S375, T376 and T379), sheep (GRK2), goat (GRK5) or mouse (HiBiT) and diluted to 1:500 (T370, S375, T376, T379 and HiBiT), 1:1,000 (pSrc-Y416, β-arrestin2, GRK2 and GRK5) or 1:2,000 (Src and GAPDH) in either BSA (T370, S375, T376, T379, HiBiT, pSrc-Y416) or 5% (w/v) non-fat dried milk powder (Src, GRK2, GRK5 and GAPDH) in TBS-T (0.1%). Horseradish peroxidase conjugated secondary antibodies against rabbit, sheep and goat were purchased from ThermoFisher (UK) and diluted to 1:10,000 in the relevant blocking buffer. IRDye conjugated secondary antibodies raised in donkey against rabbit (dye 680) and mouse (dye 800CW) were purchased from Licor (USA) and diluted in BSA (3.75%) to 1:800.

### μ receptor phosphorylation assay

μ receptor c-terminal phosphorylation was assessed using an in-cell western blot approach with established phosphorylation-specific μ receptors antibodies (Gillis et al., 2020; Underwood et al., 2024). HEK cells stably expressing HiBiT-tagged human μ receptors were plated in poly-D-lysine coated (500 μg/ml) 96-well black walled μ-clear tissue culture plates (Greiner) in culture medium at a density of 35,000 cells per well and left to settle overnight. The following day, culture medium was replaced with 100 μl of Opti-MEM containing eCF506 (1 μM) or an equal volume of DMSO and incubated overnight. Opti-MEM was replenished the following morning with 80 μl of fresh Opti-MEM before 20 μl of DAMGO (1 nM - 10 μM, prepared at 5x concentration in Opti-MEM) or DMSO was serially diluted into each well and incubated at 37°C for 20 minutes. Agonists were removed and cells immediately fixed using 100 μl paraformaldehyde (4% v/v) in tris-buffered saline (TBS) for 5 minutes at room temperature. Antigen retrieval was performed for 5 minutes at 37°C using trypsin (0.025%) in CaCl_2_ containing buffer (pH 7.8) prior to washing each well three times with TBS. Cells were permeabilised for 15 minutes using triton X-100 (0.1% v/v) followed by blocking in 5% (w/v) BSA reconstituted in TBS for 1 hour at room temperature.

Primary antibodies directed against phosphorylated μ receptor c-terminal residues (rabbit anti-pT370/pS375/pT376 or pT379) or mouse anti-HiBiT tag were co-diluted 1:500 in 3.75% BSA containing 0.1% tween-20 (TBS-T) and co-incubated (75 μl per well) overnight at 4°C. Some wells were incubated with 3.75% BSA only as a primary antibody control. The following morning, primary antibodies were removed and stored at 4°C for reuse up to three times and wells washed with TBS-T. Secondary antibodies (Licor donkey anti-rabbit 680RD and donkey anti-mouse 800 reconstituted in 3.75% BSA in TBS-T) were incubated at room temperature for 1 hour in the dark. Some wells were incubated with 3.75% BSA in TBS-T only as a secondary antibody control. Secondary antibodies were discarded, wells washed three times to remove unbound secondary antibody, and the plate dried before scanning on a Licor Odyssey Clx with a resolution of 84 μm and an offset of 3.0 mm. Fluorescence intensities in the 700 nm wavelength (phosphorylated μ receptor c-terminal residues) were expressed relative to fluorescence in the 800 nm wavelength (HiBiT-tagged total μ receptor) in each well and normalised to the percentage of maximum fluorescence for each phosphorylated residue measured in DMSO exposed cells, respectively.

### c-Src plasmids

All constructs are based on human c-Src WT (1-536) c-Src (plasmid #42202; RRID: Addgene_42202), c-terminus truncated (1-249) c-Src (plasmid #42204; RRID: Addgene_42204), n-terminus truncated (250-536) c-Src (plasmid #42208; RRID: Addgene_42208), and kinase dead (K298M) c-Src (plasmid #42210; RRID: Addgene_42210), were donated by Dr Robert Lefkowitz. Constitutively active c-Src (Y530F) (plasmid #124659; RRID: Addgene_124659) was donated by Dr Cheng-Han Yu. All plasmids were amplified as recommended using the PureYield™ Maxiprep System (#A2393; Promega, UK) in chemically competent DH5α.

## Data and statistical analysis

Concentration-response relationship data derived from functional assays of μ receptor activation were expressed as a percentage of the maximum luminescence produced by a reference agonist (DAMGO) without experimental treatment present on each plate (β-arrestin2 recruitment), or the luminescence produced by luciferin and forskolin in each well prior to agonist exposure (cAMP accumulation). These data were plotted and fitted to a three-parameter logistic function (Response = minimum + ((maximum-minimum)/1+10^(log(potency)-[agonist])^)) yielding maximum and minimum responses used to derive estimates of agonist efficacies (E_MAX_) and potencies (EC_50_ or IC_50_). The latter converted to positive logarithmic molar values (pEC_50_ or pIC_50_, M) to generate a Gaussian-distributed data set amenable to parametric analysis. Constraints to individual fits were set to minimum ≥0 for β-arrestin2 recruitment and for cAMP accumulation assays.

Assumptions of parametric analyses were confirmed prior to statistical comparisons using data derived from individual concentration-response relationships. Averaged data were also fitted with the logistics function and are shown for illustrative purposes. E_MAX_ and pEC_50_ values in assays of β-arrestin2 recruitment, or E_MAX_ and pIC_50_ values in assays of cAMP accumulation, were compared between different inhibitor and vehicle control treatments using a one-way ANOVA. E_MAX_ and pIC_50_ values from experiments involving depletion of available μ receptors were compared between their respective values in cells with full receptor availability using an unpaired T-test. C-terminal phosphorylation of HiBiT tagged human μ receptors were compared using a two-way ANOVA with concentration and overnight treatment as factors. Summary data presented are mean ± SEM (E_MAX_) or mean with upper and lower quartiles (pEC_50_/ pIC_50_) of at least 5 separate experiments unless stated in the associated legend. Reported ‘n’ values in the figure legends represent the number of individual 96-well plates. Significance indicated in the summary tables and graphs represent p < 0.05 and post-hoc pairwise comparisons with a Bonferroni or Dunnet correction applied were only performed if F achieved p < 0.05. Construction of graphs and all statistical analyse were performed using GraphPad Prism (version 9.5.1).

## Acknowledgements

We thank the staff of the University of Dundee MSRU for assistance with animal experiments and husbandry and Stella Reynolds for help with some assays. This study was supported by a NIAA (BJA/RCoA) grant WKR0-2019-0067 awarded to TGH and a Tenovus Scotland grant awarded to SS and TGH (T23-14). SS and TGH are part of the Advanced Pain Discovery Platform, supported by a UKRI and Versus Arthritis Grant: MR/W002566/1.

## Author contributions

SS, AUB and TGH conceived and designed all aspects of this study and wrote the first draft of the

manuscript and edited subsequent versions. SS analysed the data. ALM and AUB synthesised eCF506.

RT performed cloning. SS, FN, SR, EL, ALM, RT and AUB generated the data used throughout this manuscript. TGH and SS acquired funding.

## Disclosure and competing interest statement

The authors have no competing interests or disclosures.

## Data availability

The data that support the findings of this study are available from the corresponding author, upon reasonable request.

